# Detection of Diffuse to Focal Myocardial Fibrosis by Cardiovascular Magnetic Resonance Against Histology in Mini-Swine

**DOI:** 10.1101/2025.01.13.632871

**Authors:** Huaying Zhang, Cui Chen, Wenjing Yang, Di Zhou, Yining Wang, Leyi Zhu, Mengdi Jiang, Qiang Zhang, Shihua Zhao, Stefan K Piechnik, Vanessa M Ferreira, Minjie Lu

**Affiliations:** Department of Magnetic Resonance Imaging, Fuwai Hospital, State Key Laboratory of Cardiovascular Disease, National Center for Cardiovascular Diseases, Chinese Academy of Medical Sciences and Peking Union Medical College, Beijing, China; Oxford Centre for Clinical Magnetic Resonance Research (OCMR), John Radcliffe Hospital, National Institute for Health Research (NIHR) Oxford Biomedical Research Centre, Oxford BHF Centre of Research Excellence, University of Oxford, Oxford, United Kingdom; Big Data Institute, Nuffield Department of Population Health, University of Oxford; Key Laboratory of Cardiovascular Imaging (Cultivation), Chinese Academy of Medical Sciences, Beijing, China

**Keywords:** Cardiovascular Magnetic Resonance, T1-mapping, Myocardial Fibrosis, Histology, Swine

## Abstract

**Background:** Myocardial fibrosis predicts adverse outcomes in myocardial infarction (MI) and other cardiovascular conditions. Cardiovascular magnetic resonance (CMR) can non-invasively detect focal and diffuse myocardial fibrosis, but comprehensive validation against histology remains scarce.

**Aims:** To assess the comparative diagnostic performance of CMR methods in detecting myocardial fibrosis, using a mini-swine model and histology as gold standard.

**Methods:** Eighteen mini-swine (16 MI; 2 healthy) underwent CMR cine, LGE, T1- and ECV-mapping. Two commonly-used T1-mapping methods – MOLLI 5(3)3 and ShMOLLI 5(1)1(1)1 – were included. Pathological sections were categorized as infarcted, peri-infarct, remote and healthy myocardium based on triphenyl tetrazolium chloride staining. Fibrotic burden was quantified by collagen volume fraction (CVF) into severe (CVF≥30%), moderate (CVF=10-25%), and mild (CVF=3-14%). The relationships between LGE, T1, ECV and CVF, and diagnostic performance using area-under-the-curve (AUC), were analyzed.

**Results:** For detecting severe fibrosis, LGE, T1 and ECV all had excellent diagnostic performance (AUC: LGE=0.93, ECV_ShMOLLI_=0.96, T1_ShMOLLI_=0.91, T1_MOLLI_=0.93, ECV_MOLLI_=0.88). ECV_ShMOLLI_ showed significantly better discriminatory accuracy than ECV_MOLLI_ in detecting severe fibrosis and MI (both p<0.05), with the highest correlation to CVF (ECV_ShMOLLI_ r=0.86, ECV_MOLLI_ r=0.82, T1_ShMOLLI_ r=0.77, T1_MOLLI_ r=0.77, semi-quantitative LGE r=0.75). Only T1_ShMOLLI_ and ECV_ShMOLLI_, but not LGE or T1_MOLLI_/ECV_MOLLI_, differentiated remote myocardium with mild fibrosis (CVF=8.23%) from healthy myocardium (CVF=2.01%).

**Conclusions:** CMR can detect severe to mild myocardial fibrosis as validated against histology. For low-grade fibrosis, T1-mapping significantly outperformed LGE. Choice of CMR methodologies matters for myocardial fibrosis detection, which has importance for clinical trial design.

## Section 1: Introduction

Myocardial fibrosis is commonly seen in the end stages of a wide range of cardiac conditions, including both ischemic and non-ischemic heart disease^1,2^. On histopathology, myocardial fibrosis may present as focal or diffuse fibrosis^3^. Myocardial fibrosis can lead to not only myocyte contractile dysfunction, increased cardiac stiffness and heart failure, but may also serve as a substrate for arrhythmias and sudden cardiac death^4^. The presence of myocardial fibrosis has prognostic significance, often correlating to major adverse cardiac events^5^.

In acute myocardial infarction, the process of developing myocardial fibrosis is not only activated in the infarcted core, which eventually becomes a focal scar^6^, but is also involved in the progression of injury and repair in the non-infarcted remote myocardium as more subclinical diffuse fibrosis, which can lead to adverse ventricular remodeling and dysfunction^7^. While focal myocardial fibrosis is generally irreversible^8^, diffuse fibrosis, especially low to moderate grade, is potentially reversible^9^. Thus, the ability to detect both focal and diffuse fibrosis not only offers value in prognostication but also opportunities for early intervention to favorably alter patient outcomes. Accurately detecting the presence of diffuse fibrosis may also open new venues to assess the efficacy of emerging anti-fibrotic treatments^10^.

Cardiovascular magnetic resonance (CMR) is the imaging standard for detecting myocardial fibrosis non-invasively^11^. Late gadolinium enhancement (LGE) can detect focal myocardial fibrosis in both ischemic and non-ischemic heart disease^12,13^, but has limited ability to detect diffuse myocardial fibrosis^15^, where CMR methods such as quantitative pixel-wise myocardial T1-mapping, including quantification of the extracellular volume fraction (ECV), are used^16^. However, comprehensive validation of the relative ability of LGE, T1- and ECV-mapping in detecting focal and diffuse myocardial fibrosis against histopathology remains scarce.

This study aims to bridge this gap by undertaking a head-to-head comparison of CMR LGE, T1-mapping and ECV in their ability to detect a range of myocardial fibrosis in a comprehensive, prospective, histopathological study of swine MI models. Further, two commonly-used T1 methods – modified Look-Locker inversion recovery (MOLLI)^17^ and shortened MOLLI (ShMOLLI)^18^ – were directly compared, especially in their detection of low-grade myocardial fibrosis. We assessed the accuracy and correlation of LGE, native T1 mapping and ECV with collagen volume fraction (CVF) on histology as the reference gold standard, in detecting severe to mild myocardial fibrosis. This analysis is anticipated to provide clinical validated evidence for more nuanced diagnosis and management of cardiac diseases using these CMR methods, as well as have importance in clinical trial design.

## Section 2: Materials and Methods

### 2.1 Consent

Ethics approval was obtained from the Ethics Committee for Animal Study at our hospital (Fuwai Hospital) and the Care of Experimental Animals Committee of the Chinese Academy of Medical Sciences and Peking Union Medical College (0090–148-GZ [X]). The animals used in this study were reported in accordance with the ARRIVE guidelines for reporting experiments involving animals^19^. Provision of the CMR imaging technique was reviewed and approved by the University of Oxford Animal Care and Ethical Review Committee (APA/1/5/ACER).

### 2.2 Animal Model

Twenty healthy male Chinese miniature swine were prospectively included (experimental MI group, n = 18; healthy control group, n = 2; allocated randomly based on a computer based random order generator), weighing 40-48kg (average 44.8kg) at the beginning of the experiment. The age and weight of the experimental group and the healthy control group were matched.

After fasting for 12 hours, the swine were sedated with diazepam (1 mg/kg, i.m.). Anesthesia was induced with ketamine (10-15 mg/kg, i.v.). After intubation, anesthesia was maintained with 1.5–2.5% isoflurane and oxygen. Amoxicillin (15 mg/kg) and 10 mL fentanyl per swine (50 μg/mL) was administered i.m. at induction. The MI animal model was built up as previously described^20^. To put it briefly, a 3.0-mm ameroid constrictor (MRI-3.00-TI; Research Instruments NW), which can gradually shrink the blood vessels over time, was placed at the proximal end of the left anterior descending coronary artery in the experimental group. Surgical procedures in the healthy control group were the same as the experimental group except for the constrictor implantation. All surgeries were done by the same surgeon.

After the surgery, swine were maintained under anesthesia and assisted ventilation until they were hemodynamically stable and showed acceptable blood gas concentrations. Decannulation was attempted when their respiratory rate was greater than 12 breaths per min and end-tidal carbon dioxide concentration was less than 6.5%. The swine were transferred to the recovery room which was appropriately heated (around 20°C), noise-free and dimly lit. Round-the-clock nursing was given for the first 48 hours. A fentanyl patch (100 μg/h) was attached to the front leg for three days.

The swine were closely observed at least twice daily by an experienced vet throughout the rest of the study. Assessments involved post-operative pain, motor activity, changes in appearance and vital signs, etc. Coronary angiography (CAG) was performed weekly to determine the degree of coronary stenosis. The same procedure of narcosis would be repeated before CAG. The animals were included in the study if they underwent successful coronary artery occlusion, defined by a 70% or greater reduction in left anterior descending artery in the final CAG. The animals were excluded if they developed serious complications or reached humane endpoints (See Supplements Section 1).

The swine were housed separately in pens on solid concrete floors with straw bedding and wood shavings. They were fed with a commercially pelleted dry feed twice daily and water was given ad libitum. Details of animal care and welfare are provided in Supplements Section 1.

### 2.3 CMR Protocols

Both the experimental and control swine groups underwent 3T CMR (MAGNETOM Skyra, Siemens Healthcare, Erlangen, Germany) examination randomly using the same protocol four weeks (28±2 days) after the surgery and were anesthetized prior to the scan as mentioned above. Cine imaging was performed using a balanced steady-state free precession sequence in four- and two-chamber long axis as well as short-axis of the left ventricle from base to apex. Subsequently, three short axis slices (basal, middle and apical) of the left ventricle were obtained in random order (to avoid bias) for native T1 mapping, post-contrast T1 mapping and LGE images. LGE was performed using a T1-weighted phase-sensitive inversion recovery sequence about 10 minutes after injection of gadolinium-based contrast agent (GBCA) (0.2 mmol/kg; Magnevist; Bayer Healthcare Pharmaceuticals, Wayne, NJ) and post-contrast T1 mapping was acquired approximately 15-20 minutes after injection of contrast agent (Magnevist). The following sampling schemes of T1 mapping were used: native MOLLI 5(3)3 (either 256 matrix for heart rate <80 bpm or 192 matrix otherwise); pos-contrast MOLLI 4(1)3(1)2(same matrix strategy based on heart rate); native ShMOLLI 5(1)1(1)1^18,21^; post-contrast ShMOLLI 5(1)1(1)1 (Fig.1). Inline motion correction (MOCO) was applied in each sequence. Detailed imaging parameters were presented in Supplements Section 3.

**Fig. 1.**
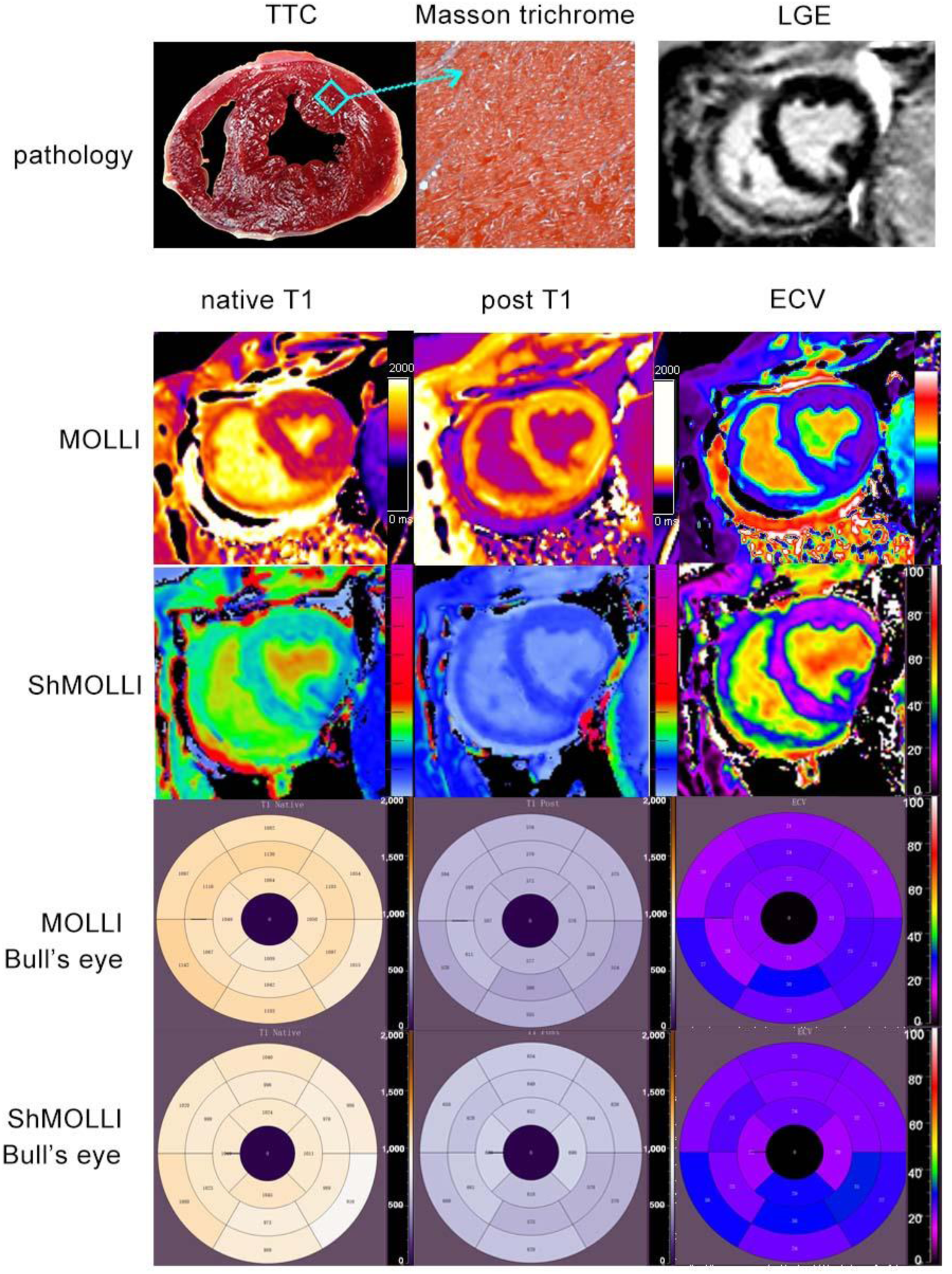
Representative mid-ventricular pathological images, LGE images and T1 mapping of the healthy control group. Pathological images of TTC staining and Masson trichrome staining as well as LGE images (first row); T1 maps of native T1, post T1 and ECV with MOLLI (second row) and ShMOLLI (third row); Bull’s eye of native T1, post T1 and ECV with MOLLI (forth row) and ShMOLLI (fifth row) Abbreviations: ECV=extracellular volume fraction; LGE=late gadolinium enhancement;TTC=triphenyl tetrazolium chloride

**Fig. 2.**
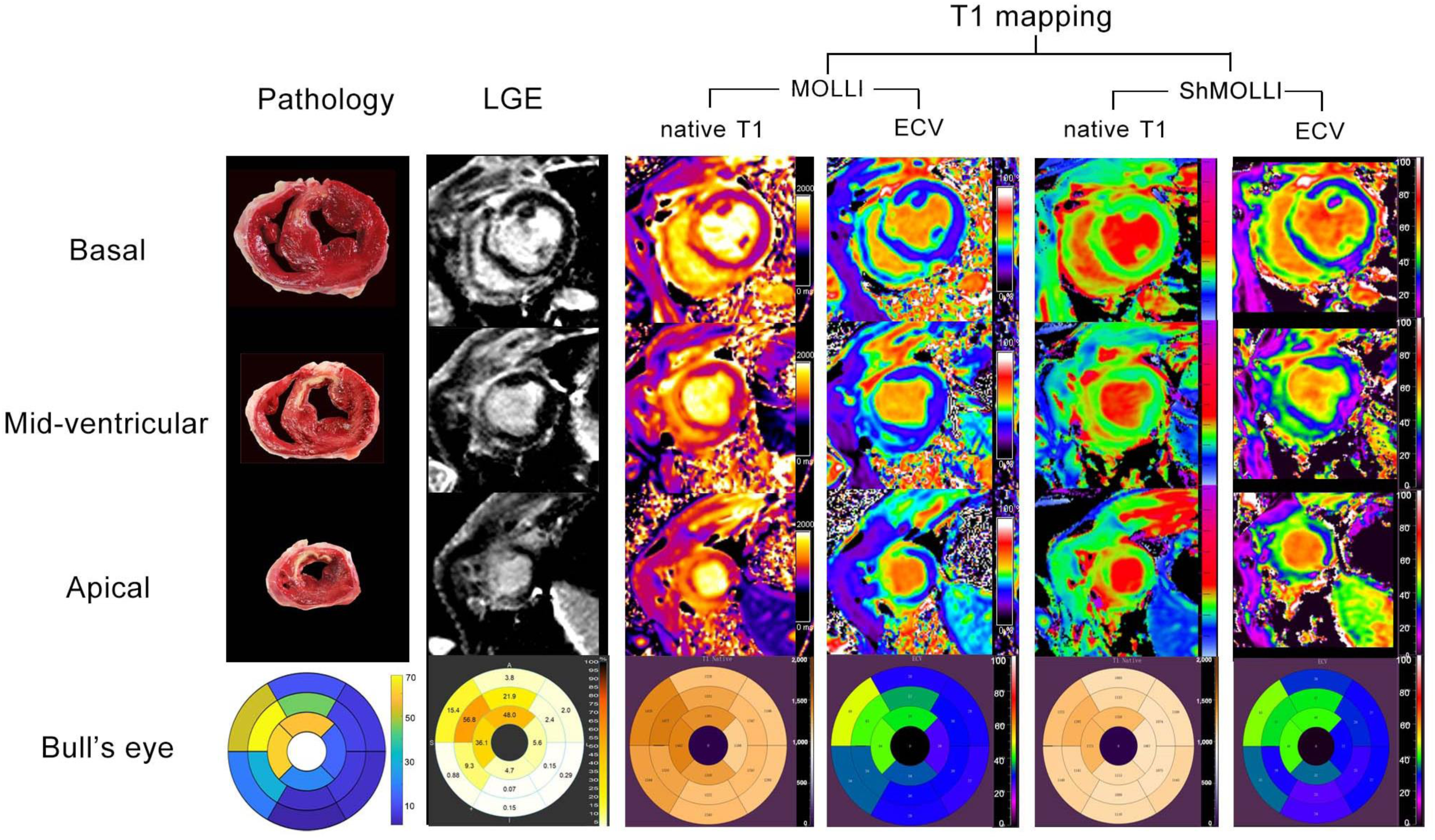
Detection of MI on pathological TTC staining, LGE images and T1 mapping. Images in a representative MI swine were shown: TTC images of basal, mid-ventricular and apical slices as well as a pathological Bull’s eye with CVF of 16 segments; LGE images, T1 mapping-MOLLI/ShMOLLI native T1 images and ECV images of the same layers as well as Bull’s eyes. Infarcted myocardium was detected at the same location on various types of images. Abbreviations: ECV=extracellular volume fraction; LGE=late gadolinium enhancement; MI=myocardial infarction; TTC= triphenyl tetrazolium chloride

### 2.4 CMR Analysis

CMR analysis was performed in a random order by using Medis Suite (Version 4.0, Leiden, the Netherlands) by two radiologists (J.X. with 10 years of CMR experience; L.Z, with 6 years of CMR experience) blinded to histopathological findings. The presence of LGE was defined as an area of LV myocardium with an increased signal intensity of ≥6 standard deviations above the mean SI of remote myocardium in the same short-axis slice of LGE image^22^; LGE burden was further quantified by the ratio of total LGE mass to the total LV mass. The T1 maps and ECV measurements were analyzed according to the American Heart Association myocardial 17-segment model with exclusion of the apical segment. For segments confounded by both infarcted and peri-infarct tissue types, delineation of regions of interest (ROI) and regions of avoidance was performed manually. ECV was calculated from pre-contrast and post-contrast T1 maps, adjusted to the hematocrit level (Hct), using the following formula: ECV = (1-Hct)×(ΔR_1_myocardium)/ (ΔR_1_bloodpool) before and after GBCA administration, where R_1_ = 1/T1^23^. Venous blood immediately obtained prior to each CMR examination was used to determine the Hct. The ECV maps provide quantitative pixel maps of ECV ranging from 0% to 100%. Representative CMR images and post-processing images of a healthy control and MI swine are displayed in Fig.1 and 2.

### 2.5 Euthanasia and Histological Validation

After the CMR examinations, the swine were maintained under isoflurane anesthesia and then humanely euthanized to obtain heart specimens. Methods of euthanasia were based on guidelines from the American Veterinary Medical Association 2020^24^. Specifically, high dose of 20 ml of 15% (volume per weight) potassium chloride was injected into the anesthetized swine intravenously rapidly, resulting in transient hyperkalemia, leading to cardiac arrest in 15-20 seconds.

The heart was cut along the short axis from base to apex by a pathologist (X.D., with 10 years of experience). The slices were then immersed in a 1% triphenyl tetrazolium chloride (TTC) solution in saline at 37°C for 10 min, and digitally photographed (pathological images in Fig.1 and 2).

As is illustrated in Fig.3, pathologic slices were analyzed based on segmental level and matched with the corresponding CMR images utilizing landmarks such as papillary muscles and right ventricular insertion points. Each segment was considered to be an experimental unit. This process was performed by a pathologist (X.D.) and a radiologist (J.X.) by consensus. In the experimental swine group, the myocardial segments were categorized as infarcted, per-infarcted, and remote segments (Fig.4). An infarcted segment was defined as an area with negative TTC staining involving more than 50% of the wall thickness. A segment that was next to the infarction or had a negative TTC staining of less than 50% was classified as a peri-infarct segment. A segment contralateral to the infarcted segment with positive TTC staining and normal regional wall motion was considered as a remote segment. The sections within the normal control group were categorized as healthy control segments and defined as: areas with normal CMR images, positive TTC staining and without upstream angiographic coronary stenosis or abnormal regional wall motion.

**Fig. 3.**
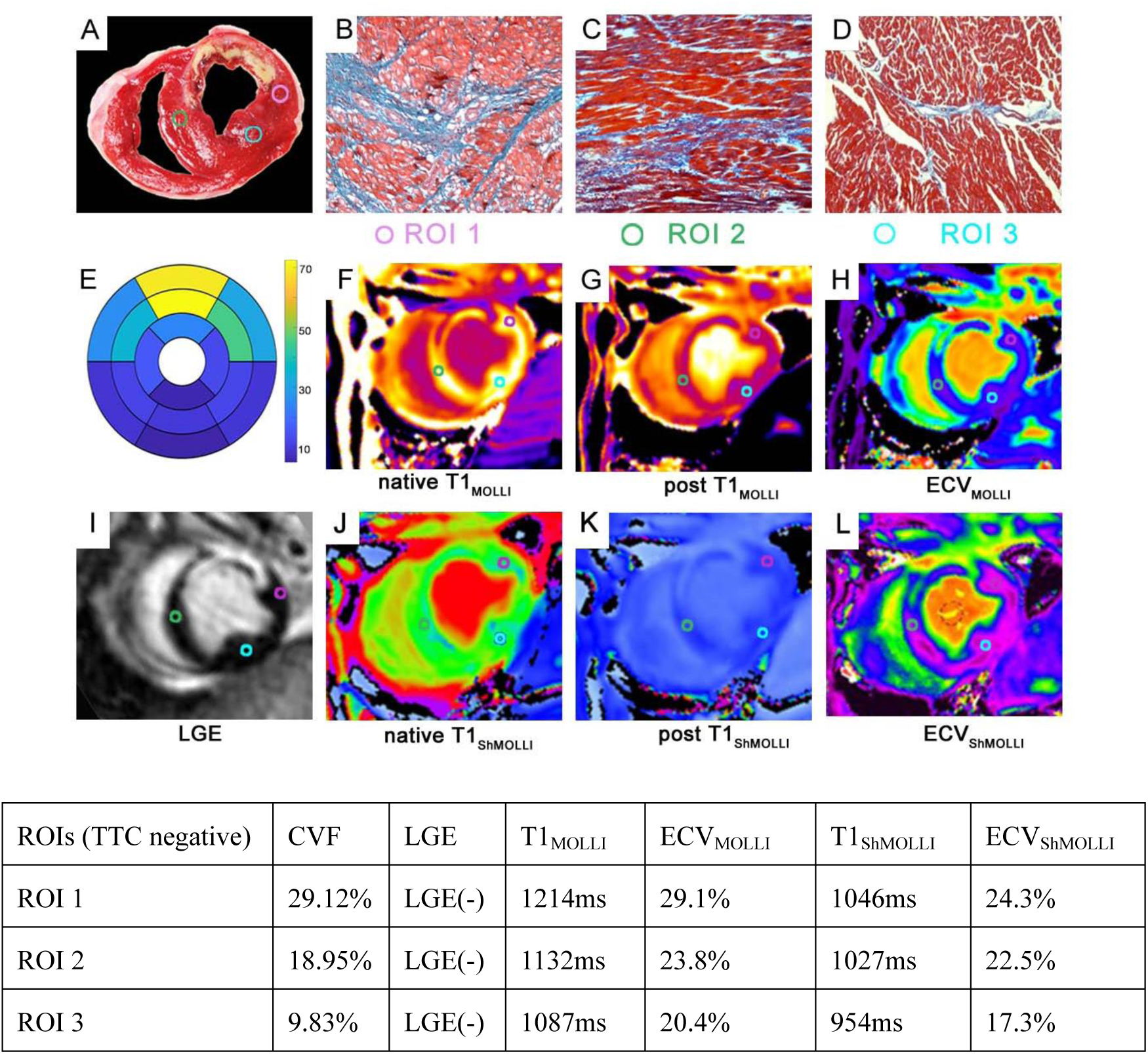
Examples of matching process between pathology and CMR imaging. A sample (ROI 1, pink circle) was taken from anterolateral wall of the basal slice (A) and calculated histologic CVF under a microscope (B). Corresponding CMR information was assessed at the same location (segment #6 in the 17-segment model) in LGE (I), MOLLI T1-maps (F-H), and ShMOLLI T1-maps (J-L). The same procedure was applied for ROI 2 and ROI 3. ROIs in the same position were marked with circles of the same color. (A)The cardiac specimen with TTC; (B-D)ROI samples with Masson trichrome; (E)Historical CVF bull’s eye; (F-H)MOLLI T1-maps with ROIs; (I)LGE imaging; (J-L)ShMOLLI T1-maps with ROIs. Abbreviations: CMR=cardiac magnetic resonance; CVF=collagen volume fraction; LGE= late gadolinium enhancement; ROI=regions of interest; TTC= triphenyl tetrazolium chloride

**Fig. 4.**
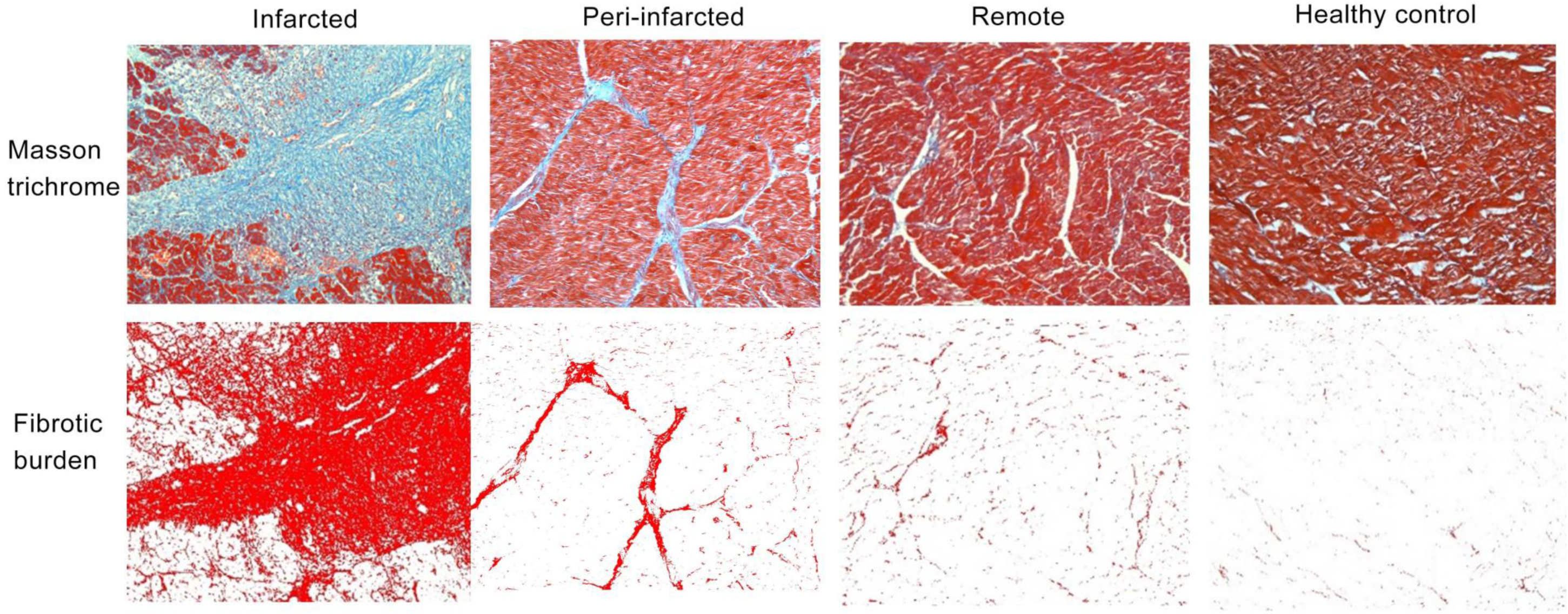
Histopathological images of representative myocardial tissue types in swine. Masson trichrome staining of pathological samples from MI swine and healthy controls (first row); software analyzes for fibrotic burden by ImageJ (second row). The fibrotic burden of representative fields of infarcted, peri-infarct, remote and healthy control segment were: CVF 67.70%, 14.11%, 1.51% and 0.59%, respectively. Abbreviations: CVF= collagen volume fraction; MI=myocardial infarction

Morphological changes of myocardial tissue were assessed by Masson trichrome staining as previously described^25^. High-power magnification (×200) digital images for 16 segments of each animal were acquired to analyze CVF. On average, 12 inconsecutive high-power fields were assessed per segment. Semiautomatic image analysis using software (ImageJ, version 1.8.0; National Institutes of Health) was performed in a cross-stitch manner. The area of collagen was determined as a percentage of the total area of the myocardium by combining the SD of the average signal and the isodata automatic threshold^26^ (Fig.4). Segments that cannot be correctly identified by TTC staining or analyzed by the software were excluded.

### 2.6 Statistical Analysis

Analyses were conducted based on segments. Normality of data was assessed using the Shapiro-Wilk test. Continuous variables were given as means ± SDs for normally distributed data or as median values with interquartile range, otherwise. Comparisons between variables of different kinds of myocardium (infarcted, peri-infarct, remote and control) were performed using one-way ANOVA or nonparametric tests. A paired t-test was employed to determine the differences among LGE, T1-map and ECV, and between MOLLI and ShMOLLI T1-mapping methods. Two-side p<0.05 was considered statistically significant. The associations of MOLLI/ShMOLLI T1 mapping parameters and CVF were assessed with *Pearson* correlation and univariable general linear regression models.

Receiver operating characteristic (ROC) curves were built to evaluate the accuracy of native T1 and ECV in detecting the existence of fibrosis using pathologic findings as the standard of reference. Specifically, in terms of the diagnostic ability of infarcted core, severe fibrosis was defined as a segment with CVF≥30%^27^ and was assigned a value of 1, whereas an absence (CVF<30%) was given a value of 0. Infarction, which was assigned a value of 1, was defined by TTC as described above, and the rest were assigned a value of 0. Further, regarding differential ability of low-grade fibrosis, segment with CVF<30% (except for controls) or remote segment was assigned a value of 1 and healthy control segment was assigned a value of 0. The optimal cutoff values were determined by Youden index. Area-under-the-curve (AUC), diagnostic thresholds, sensitivity (SN) and specificity (SP) were reported.

All statistical analyzes were conducted using IBM SPSS Statistic for Windows (version 23.0), OriginPro 2021 (version 9.8.0.200) and MedCalc software (version 22.18).

## Section 3: Results

All surgical procedures were successfully completed. One swine died of infection 6 days post-surgery, and one died of postoperative ventricular fibrillation 2 weeks post-surgery. Both were from the MI group (death rate of 10%). The rest of the swine in the MI group developed chronic coronary artery occlusion, providing a total of 18 swine for analysis (16 in the MI group and 2 in the healthy control group) (Supplementary Table 1). All 18 swine underwent successful serial CMR scanning before euthanasia. Altogether 61 segments were excluded for poor image quality due to: motion artifacts (21 segments, 7.3%), mis-triggering (12 segments, 4.2%)] or unsatisfactory staining (28 segments, 9.7%)].

### 3.1 Myocardial Tissue Characteristics and Subgroup Analyzes

In all, 227 LV myocardial segments were eligible for histopathologic analysis, including 61 infarcted, 60 peri-infarct, 65 remote and 41 healthy control segments (Fig.4). Histological CVF and CMR imaging parameters of all 227 segments and subgroup analyzes according to the 4 tissue types are presented in Table 1.

**Table 1.** Myocardial Tissue Characteristics.

|  | All | Subgroup |  |  |  |
| --- | --- | --- | --- | --- | --- |
|  |  | infarcted | peri-infarct | remote | healthy control |
| **Number of segments (n)** | 227 | 61 | 60 | 65 | 41 |
| **LGE presence, n (%)** | 59 (26) | 53 (87)<sup>bcd</sup> | 6 (10)<sup>a</sup> | 0 (0)<sup>a</sup> | 0 (0)<sup>a</sup> |
| **native T1<sub>MOLLI</sub> (ms)** | 1124(95) | 1208±55<sup>bcd</sup> | 1140±42<sup>acd</sup> | 1109(16)<sup>ab</sup> | 1085±38<sup>ab</sup> |
| **ECV<sub>MOLLI</sub> (%)** | 24.6(5.9) | 33.5±8.1<sup>bcd</sup> | 24.8±3.1<sup>ad</sup> | 23.1(4.3)<sup>a</sup> | 22.8±2.8<sup>ab</sup> |
| **post T1<sub>MOLLI</sub> (ms)** | 602(66) | 540±71<sup>bcd</sup> | 602±47<sup>ad</sup> | 604±44<sup>ad</sup> | 637±50<sup>abc</sup> |
| **native T1<sub>ShMOLLI</sub> (ms)** | 1004(75) | 1113±79<sup>bcd</sup> | 992±48<sup>ad</sup> | 994(64)<sup>ad</sup> | 947±64<sup>abc</sup> |
| **ECV<sub>ShMOLLI</sub> (%)** | 22.3(7.0) | 32.7±7.4<sup>bcd</sup> | 21.0±1.9<sup>ad</sup> | 22.0±3.5<sup>ad</sup> | 20.1(4.0)<sup>abc</sup> |
| **post T1<sub>ShMOLLI</sub> (ms)** | 633(79) | 572±68<sup>bcd</sup> | 619±65<sup>acd</sup> | 650±71<sup>ab</sup> | 655±77<sup>ab</sup> |
| **CVF (%)** | 11.7(24.74) | 57.26±24.62<sup>bcd</sup> | 17.04±9.13<sup>acd</sup> | 8.23(6.94)<sup>abd</sup> | 2.01(2.28)<sup>abc</sup> |

LGE, T1 mapping parameters (MOLLI and ShMOLLI) and CVF of 227 segments and subgroup analyzes. According to normality tests, variables were given as means ± SDs or as median values (interquartile range) and comparisons were performed using one-way ANOVA or nonparametric tests.
a: vs. infarct segments, $p < 0.05$ ; b: vs. peri-infarct segments, $p < 0.05$ ; c: vs. remote segments, $p < 0.05$ ; d: vs. control segments, $p < 0.05$

Histological CVF demonstrated the strongest ability to differentiate among the four tissue types (all p<0.05). The burden of fibrosis in infarcted, peri-infarct, remote and healthy myocardial segments were 57.26%, 17.04%, 8.23% and 2.01%, respectively. As expected, LGE had a high positive rate (87%) in infarcted segments, a low positive rate (10%) in peri-infarct segments, and was absent in remote and healthy control segments.

Infarcted segments, as defined by TTC staining on pathology, showed a significantly higher amount of fibrosis (histological CVF=57.26%), LGE, T1 values and ECV, compared to the other 3 tissue types (all p<0.05). Peri-infarct segments showed significantly higher amount of fibrosis compared to both remote and healthy myocardium (histological CVF 17.04% vs 8.23% vs 2.01%, respectively; all p<0.05), and significantly higher T1 and ECV compared to healthy control myocardium (all p<0.05%) (Fig.3). Remote myocardium showed no LGE but mild degrees of fibrosis on histological CVF compared to healthy myocardium (CVF 8.23% vs 2.01%, respectively; p<0.05). Only histological CVF, T1_ShMOLLI_ and ECV_ShMOLLI_ were able to differentiate remote myocardium from healthy control myocardium, whereas LGE and MOLLI-based T1 and ECV did not differentiate the two.

### 3.2 Relationship among T1, ECV, and CVF

Across the spectrum of myocardial fibrosis, semi-quantitative LGE was highly correlated to histological CVF (LGE r=0.75, p<0.05) (Fig. 5E). ECV showed significantly better association with CVF than LGE and native T1 (ECV_ShMOLLI_ r=0.86, ECV_MOLLI_ r=0.82, T1_ShMOLLI_ r=0.77, T1_MOLLI_ r=0.77; all p<0.05) (Fig. 5A-D). ECV_ShMOLLI_ had a trend level of better correlation with CVF than ECV_MOLLI_ (r=0.86 vs. r=0.82, p=0.076). There were moderate correlations between the two T1-mapping methods tested (native T1 r=0.55, ECV r=0.70, both p<0.01).

**Fig. 5.**
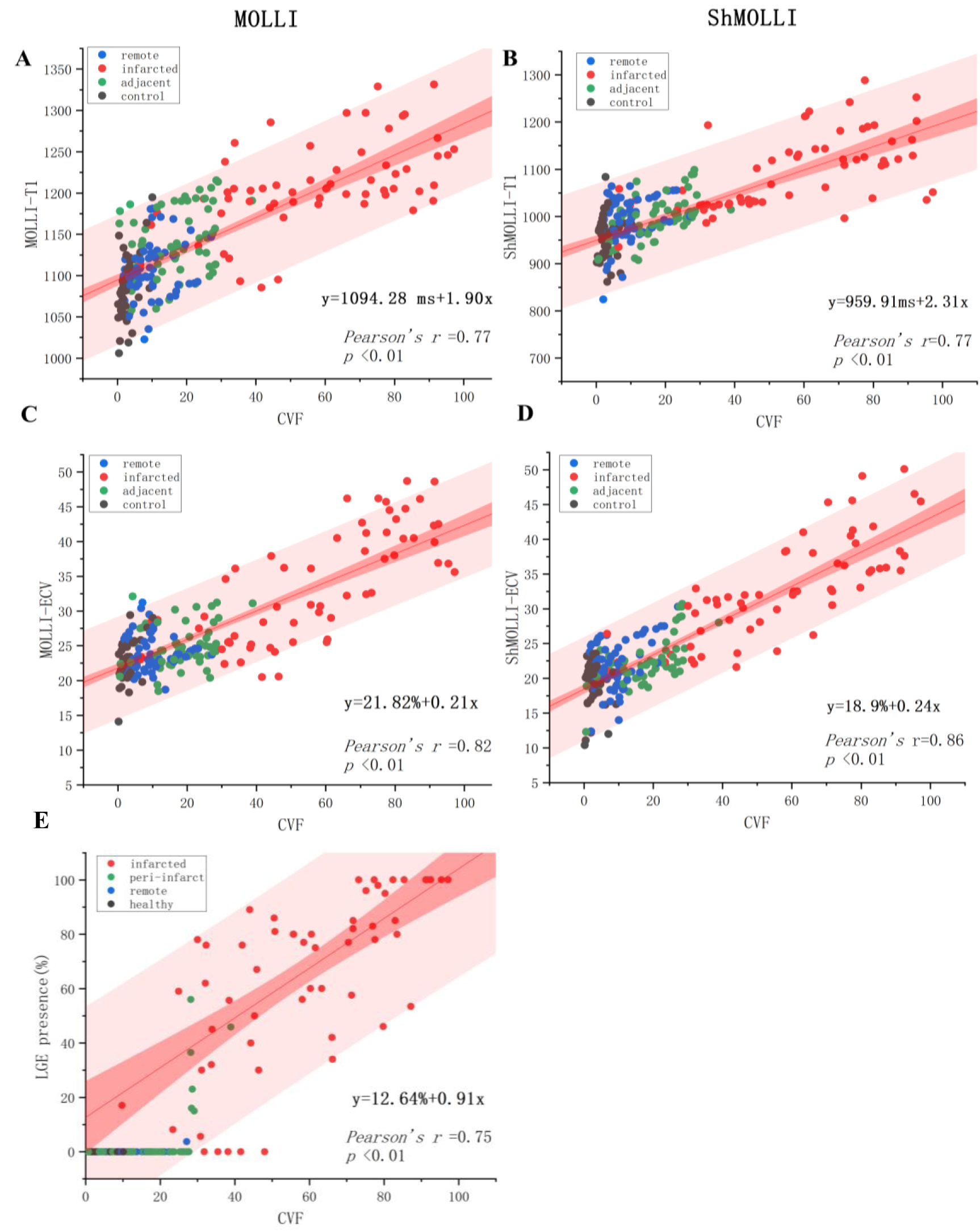
Linear regressions of T1 mapping parameters and LGE presence against histological CVF. Colored dots represented different tissue types (red dots for infarcted segments, green dots for peri-infarct segments, blue dots for remote segments and black dots for healthy segments). Red band represented 95% confidence interval and pink band represented 95% prediction band. The linear regression equations and *Pearson* correlation r were presented in bottom left corner of each figure. A. Linear regressions of native T1_MOLLI_ against CVF; B. Linear regressions of native T1_ShMOLLI_ against CVF; C. Linear regressions of ECV_MOLLI_ against CVF; D. Linear regressions of ECV_ShMOLLI_ against CVF; E. Semi-quantitative LGE presence against CVF. Abbreviations: CVF=collagen volume fraction; ECV=extracellular volume fraction; LGE=late gadolinium enhancement

Of note, in remote segments, there was mild fibrosis, as demonstrated in histological CVF (3%-14%), but both TTC staining on gross pathology and LGE were often negative (Fig 3). In remote segments, the ShMOLLI T1-mapping method showed significant correlation to histological CVF (ECV_ShMOLLI_ r=0.33, T1_ShMOLLI_ r=0.41 in remote segments; both p<0.05), in contrast to the MOLLI T1-mapping method (ECV_MOLLI_ r=0.02, T1_MOLLI_ r=0.24 in remote segments; both p>0.05), indicating better diagnostic value of the ShMOLLI T1-mapping method in detecting mild fibrosis (histological CVF 3%-14%).

### 3.3 Diagnostic Performance of CMR methods in detecting mild to severe myocardial fibrosis

For detection of severe myocardial fibrosis (defined by CVF on histology) or MI (defined by TTC staining on pathology), all CMR methods (LGE, T1 and ECV) showed good diagnostic performance (AUC 0.87-0.96; Fig.6A&B). ECV_ShMOLLI_ demonstrated the best diagnostic ability in detecting both severe fibrosis (AUC=0.96; p<0.05) and MI (AUC=0.93; p<0.05), and outperformed ECV_MOLLI_ (AUC 0.87-0.88; both comparisons p<0.05). T1_ShMOLLI_ and T1_MOLLI_ had similar performances in detecting severe fibrosis and MI (AUC 0.90-0.93; both comparisons p>0.05). There were no differences between LGE and T1-mapping parameters (AUC 0.91-0.93; all p>0.05).

**Fig. 6.**
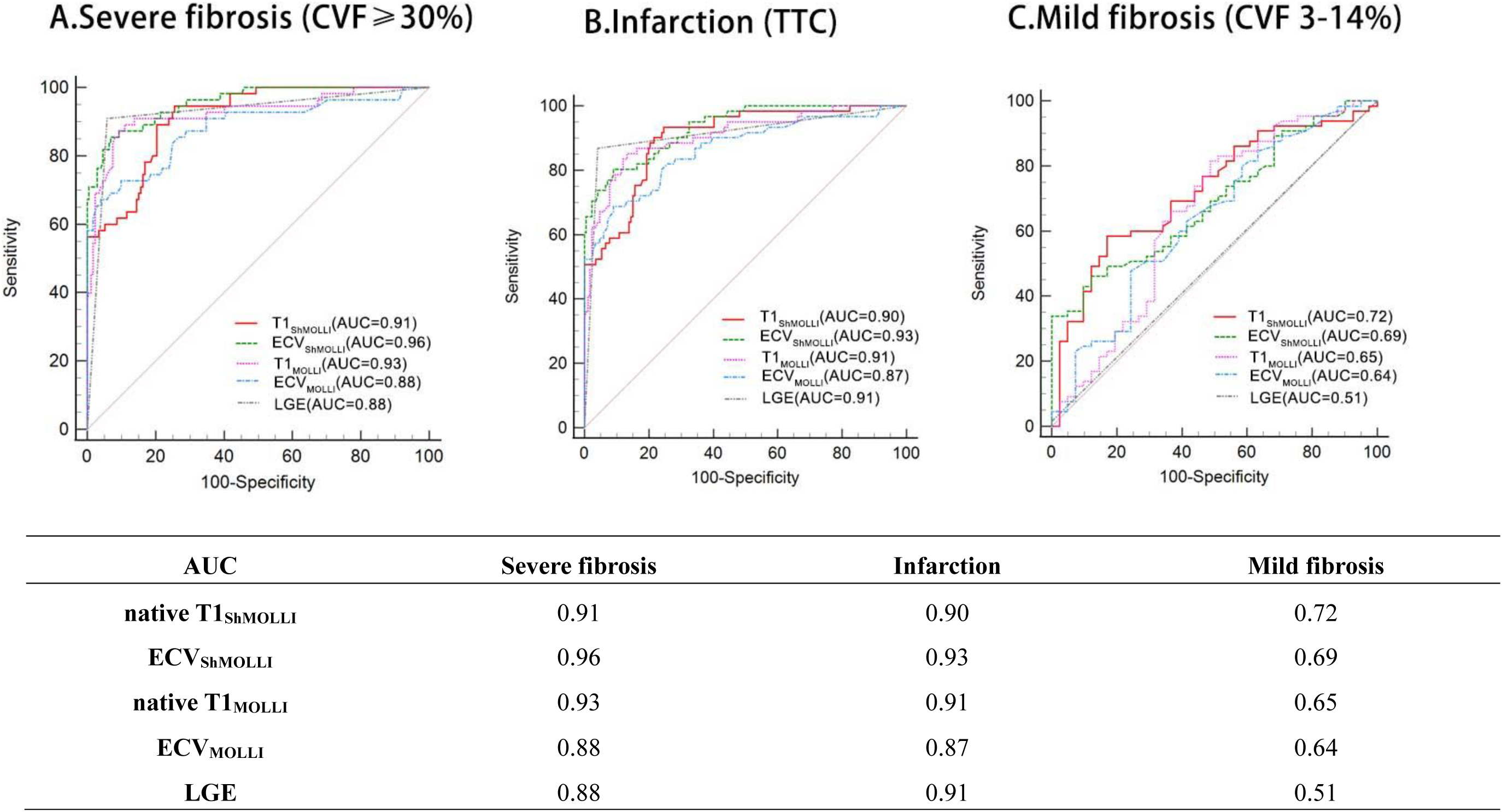
Relative diagnostic performance of CMR LGE, T1 and ECV in detecting myocardial fibrosis versus healthy normal myocardium in swine. (A) Severe myocardial fibrosis (histological CVF≥30%); (B) MI (by TTC staining on gross pathology); (C) Mild fibrosis (remote myocardium with histological CVF IQR 3-14%). Details of each ROC curve including *p*-value, sensitivity, specificity and cut-off value can be found in Supplementary Table 2. Abbreviations: CVF=collagen volume fraction; ECV=extracellular volume fraction; LGE=late gadolinium enhancement; ROC=receiver operating characteristic; TTC= triphenyl tetrazolium chloride

For detection of mild myocardial fibrosis (such as CVF 3-14% in remote segments), T1 mapping demonstrated clear advantages over LGE (all p<0.05). T1_ShMOLLI_ performed best in detecting mid myocardial fibrosis (AUC=0.72 vs AUC=0.65 for T1_MOLLI_), but statistical differences between T1-mapping parameters were not found (Fig. 6C).

## Section 4: Discussion

Our study provided a comprehensive histopathological validation of commonly-used CMR methods for detecting the spectrum of myocardial fibrosis in a swine model of myocardial infarction (MI). For detecting severe myocardial fibrosis, LGE, T1 and ECV all had excellent diagnostic performance, with ECV (using the ShMOLLI T1-mapping method) showing significantly better discriminatory accuracy than others and with the highest correlation to CVF. Additionally, in areas with low-grade fibrosis, which TTC staining and LGE was unable to detect, low-grade collagen deposition was validated by historical CVF and can be detected by T1 mapping. Furthermore, the performance of the ShMOLLI T1-mapping method was superior to MOLLI T1-mapping method in detecting mild fibrosis (CVF=8.23%) in remote myocardium from healthy myocardium (CVF=2.01%). These insights are valuable for aiding to select the most appropriate CMR methods for detecting a range of myocardial fibrosis in clinical practice and in clinical trials.

To our knowledge, this is the first head-to-head comparison among CMR imaging methods for fibrosis detection techniques, using macro cardiac specimens and micro historical CVF as reference. TTC staining is the gold-standard for the definition of infarction^28^ and LGE is the routine CMR method for detection myocardial scarring, such as from MI^29^. By segmenting myocardial tissue into different regions, we demonstrated T1 mapping was more sensitive and accurate in identifying fibrosis than TTC and LGE, and introduced a more comprehensive understanding of various myocardial response to MI. In addition to extensive necrosis and fibrosis in the core of the infarction, which TTC staining and LGE can detect in most cases, diffuse myocardial fibrosis was also detected in both the peri-infarct and remote myocardium post-MI by microscopic pathology, which showed increased native T1 and ECV in CMR. In other words, T1 mapping can simultaneously identify focal myocardial scarring and diffuse myocardial fibrosis in an efficient, single examination.

In previous studies of MI patients undergoing T1 mapping in the acute phase and follow-up (5-6 months), native T1^30^ and ECV^31^ were found to be elevated in the remote myocardium during the acute MI, and remained elevated in patients with adverse remodeling, compared to patients without T1/ECV elevation and adverse remodeling. Carrick et al^32^ showed that increased native T1 in remote myocardium was independently associated with adverse remodeling. Garg et al^33^ found that normal myocardial segments with substantially increased ECV from the acute to chronic phase showed deterioration in wall thickening and contractile function at follow-up. Shanmuganathan et al^34^ showed that high T1 in the non-infarcted myocardium in acute STEMI patients had increased risk of long-term major adverse cardiac events (2.53 [IQR: 1.03-6.22]), compared with patients with normal T1 in the non-infarcted myocardium; this added value was beyond conventional markers such as LVEF, infarct size, and microvascular obstruction. We have also, in this present study, demonstrated that the presence of increased CVF in non-infarcted remote myocardium and the superior diagnostic value of T1 mapping in detecting low-grade myocardial fibrosis. Diffuse fibrosis may start early post-MI and play a role in adverse ventricular remodeling in the longer term.

Briefly, from an CMR methodological perspective, both the ShMOLLI and MOLLI T1-mapping methods produce high quality T1 map data with good precision and reproducibility^35,36^ and are widely used in both scientific and clinical practice^37,38^. Rapid and accurate assessment of myocardial fibrosis is particularly essential for acute MI, where ShMOLLI sequence, with its short acquisition time and conditional reconstruction algorithm^18^, significantly eliminates heart-rate sensitivity and adapts to clinical demands. Moreover, our study has demonstrated that the ECV_ShMOLLI_ has the strongest correlation with histology, with the best discriminatory accuracy for severe fibrosis and superiority in differentiating low-grade myocardial fibrosis from healthy myocardium. Its better diagnostic performance for infarcted core will translate to a 60%-70% lower requirement for sample size calculation against the MOLLI T1-mapping method, and thus with potential cost benefit in setting up clinical studies and trials^39^. Future studies should further explore the application of these CMR methods in various patient populations and assess their potential value in improving patient outcomes.

There are some limitations in this study. this study was carried out in single center using a single MR system vendor and single field strength of CMR, to ensure consistency and comparability among data; different CMR methods used under different scanning conditions will need to be assessed independently. The peri-infarct myocardium is more challenging to analyze, due to a mixture of necrotic, edematous and fibrotic tissue, all of which can confound T1 elevation, and thus not used to represent a myocardial fibrosis tissue class. Despite the great fidelity of the MI swine model in representing chronic myocardial infarction, there may still be potential systematic biases between animal models and human studies.

## Section 5: Conclusions

CMR can detect severe to mild myocardial fibrosis in high agreement with histology. For severe fibrosis, LGE, T1 and ECV of CMR all have excellent diagnostic performance. For low-grade fibrosis, T1-mapping provides superior performance compared to LGE. Choice of CMR methodologies matters for the sensitive and accurate detection of myocardial fibrosis, which has importance for more efficient clinical trial design and advanced therapies.

## Abbreviations List

AUC: area-under-the-curve

CAG: Coronary angiography

CMR: cardiovascular magnetic resonance

CVF: collagen volume fraction

ECV: extracellular volume fraction

LGE: late gadolinium enhancement

MI: myocardial infarction

MOLLI: modified Look-Locker inversion recovery

ROC: Receiver operating characteristic

ShMOLLI: shortened MOLLI

TTC: triphenyl tetrazolium chloride

## Funding and Acknowledgements

This work was supported by National Natural Science Foundation of China [grant number 81971588]; Construction Research Project of Key Laboratory (Cultivation) of Chinese Academy of Medical Sciences [grant number 2019PT310025]; Youth Key Program of High-level Hospital Clinical Research [grant number 2022-GSP-QZ-5]; National Foreign Expert Talent Project [grant numbers G2021194020L]; Undergraduate Education Reform Project of Peking Union Medical College [grant number 2023zlgl 026]; and National High-Level Hospital Clinical Research Funding(2022-GSP-QN-17). We gratefully thank Dr.Vanessa Ferreira, Dr.Stefan Piechnik and Dr.Qiang Zhang for their valuable guidance and contributions to this research. Vanessa Ferreira is the Kusuma Senior Research Fellow in Cardiovascular Medicine at the University of Oxford, and acknowledges funding from The Kusuma Trust UK.

## Central Illustration

Eighteen mini-swine underwent CMR cine, LGE, and T1 mapping (including MOLLI and ShMOLLI). Historical validation was based on TTC staining and CVF. The correlation between LGE, native T1, ECV and CVF, and diagnostic performance using ROC curves were analyzed. Abbreviations: CMR=cardiac magnetic resonance; CVF=collagen volume fraction; ECV=extracellular volume fraction; LGE=late gadolinium enhancement; ROC=receiver operating characteristic; TTC= triphenyl tetrazolium chloride

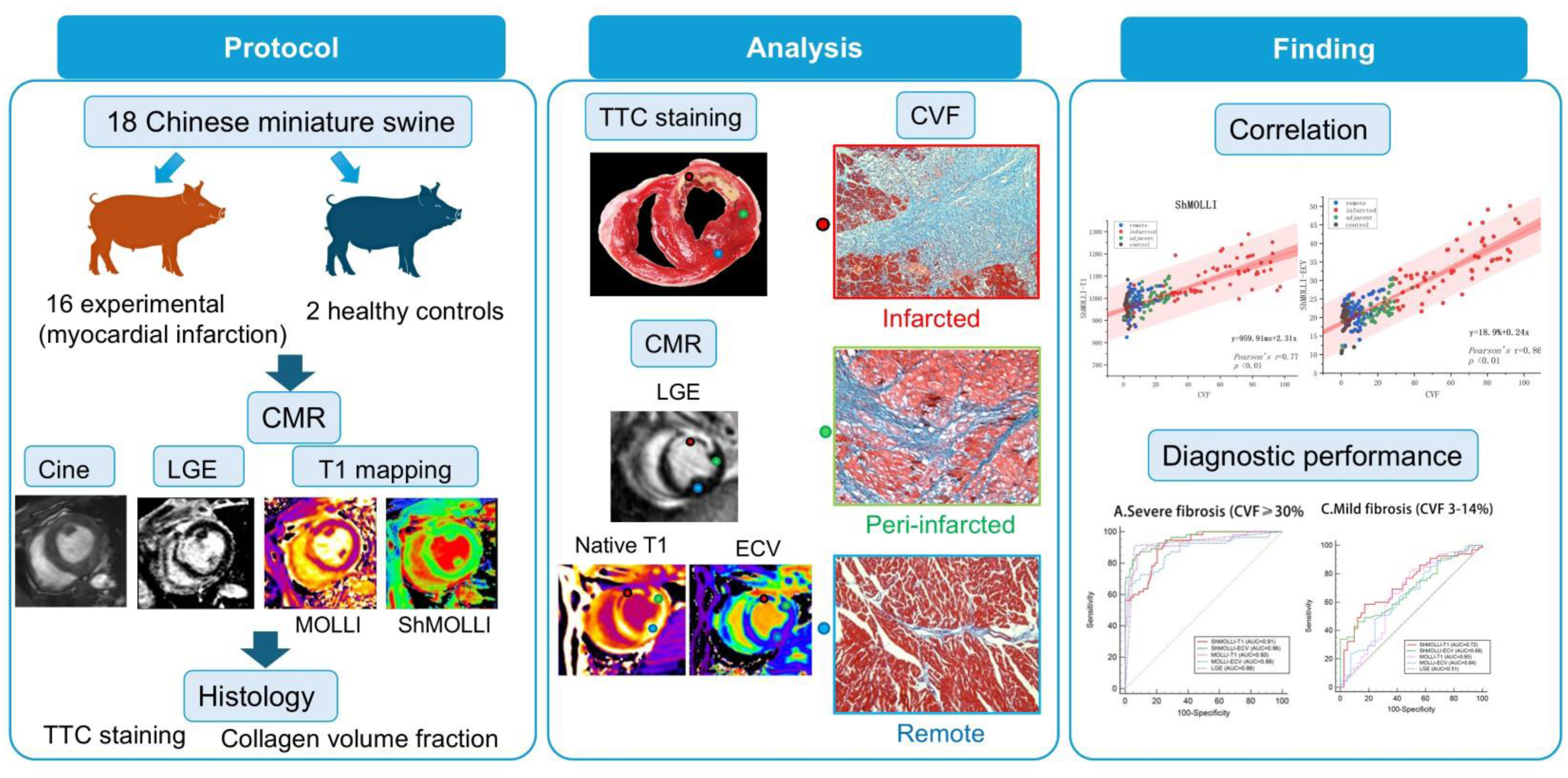

## Notes

### Competing Interest Statement

The authors have declared no competing interest.

